# BayesFM: a software program to fine-map multiple causative variants in GWAS identified risk loci

**DOI:** 10.1101/067801

**Authors:** Ming Fang, Michel Georges

## Abstract

We herein describe a new method to fine-map GWAS-identified risk loci based on the Bayesian Least Absolute Shrinkage Selection Operator (LASSO) combined with a Monte Carlo Markov Chain (MCMC) approach, and corresponding software package (BayesFM). We characterize the performances of BayesFM using simulated data, showing that it outperforms standard forward selection both in terms of sensitivity and specificity. We apply the method to the *NOD2* locus, a well-established risk locus for Crohn’s disease, in which we identify 13 putative independent signals.

## Introduction

Thousands of risk loci have been identified by Genome Wide Association Studies (GWAS) affecting nearly all studied common complex diseases in humans (Welter et al., 2014, and http://www.ebi.ac.uk/gwas/). However, for the vast majority of risk loci the causative variants and genes remain unknown. This knowledge is essential to fully capitalize on the investments in GWAS, including for the development of improved diagnostic and therapeutic applications.

A number of issues complicate the identification of the causative variants by association analysis. The first is that the utilized case-control cohorts are usually genotyped for only a subset of the variants segregating in the population. Ideally, fine-mapping would require sequencing of all cases and controls in the chromosome regions of interest, if not the entire genome. This will remain difficult to achieve, at least in the short term. At present, the best alternative is genotype imputation using for instance the data from the 1,000 Genomes Project as reference set. However, the reliability of the imputed genotypes is not perfect, particularly for low frequency and rare variants. Thus the information content varies between variants. In other words, the effective number of genotypes may vary between variants, precluding fair comparison of the strength of their association.

A second issue is the difficulty, when using the most commonly applied single-marker analyses (i.e. testing for disease association one marker at a time), to distinguish the association patterns of the causative variants from that of “passenger” variants that are merely in linkage disequilibrium (LD) with causative variants. In the case of allelic homogeneity (one causative variant only), one “asymptotically” expects the causative variant to show the strongest single-marker association (highest –log(p) value) of all variants. But in the real world the causative variant may be overshadowed by passenger variants that by chance (or as a result of unaccounted confounding effects), and given the limited size of the case-control cohorts, appear more strongly associated with the disease. The situation becomes even trickier in the case of allelic heterogeneity (i.e. the segregation of multiple causative risk variants that may or may not be in LD), a scenario that is likely to be very common. In this case, the lead SNP in single-marker analysis may be a “ghost” variant that is in LD with two or more causative variants, hence being “asymptotically” more strongly associated with the disease than either of them. Also, in the case of allelic heterogeneity, causative variants are by definition bound to exist amongst the “non lead” variants in single-marker analysis.

Improving the mapping resolution by analysis of association requires the development of statistical models that allow inclusion of confounding factors, estimation of the effects of individual variants conditional on the other ones (i.e. multi-marker analysis to distinguish causative from passenger variants), and identification of the best amongst the large number of possible models (i.e. what combination of variants, assumed to be causative, explains the data best).

We herein introduce a software package (BayesFM) that uses Bayesian Least Absolute Shrinkage Selection Operator (LASSO) combined with a Monte Carlo Markov Chain (MCMC) to achieve that goal. After describing the underlying algorithms, we test BayesFM on simulated data and compare its performances with that of a more standard “forward selection” approach. We then describe the results obtained when applying BayesFM to the *NOD2* locus, a well established risk locus for Inflammatory Bowel Disease (f.i. Jostins et al., 2015; Huang et al., 2016).

## Materials & Methods

### BayesFM algorithm

#### Assumptions

We assume that GWAS studies have identified one or more risk loci for a disease of interest in an available case-control cohort. We further assume that – within the identified risk loci - the genotypes of array-interrogated SNPs have been augmented in cases and controls with genotypes of as many variants as possible either by imputation or by sequencing. We herein propose an approach that will model disease outcome as a function of the genotypes at one or more variants in a given risk locus with the aim to fine map that locus, i.e. to identify causative variants within that locus. Fine-mapping is conducted one risk locus at the time. Risk loci defined by GWAS typically span ~250 Kb, contain ~5 genes (range: 0 to >50) and encompass thousands of common and low frequency variants.

#### Model

The proposed model is based on the standard assumption of an underlying, normally distributed liability *y* with threshold t, such that individuals for which *y > t* are affected and individuals for which *y_i_ ≤ t* are healthy. We model the liability of individual *i* (*y*) as

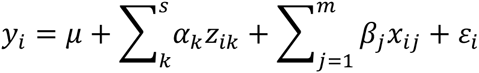

where *μ* is the population mean, *α_k_* is the effect of principal component (PC) *k* of *s, z_ik_* is the value of PC *k* for individual *i*, is the effect of variant *j* of *m, x_tj_* is the dosage of the alternative allele of variant *j* for individual *i*, and *ε_i_* is the residual error term for individual *i*. PCs were included to correct for population stratification, and values of *z_ik_* were computed using standard procedures. Other fixed effects could be added to the model in exactly the same way as PCs. *m*, i.e. the number of causative variants in the risk locus, was arbitrarily set at 20, meaning that we did not anticipate more than 20 independent effects per risk region. In other words, we consider that there can be multiple causative variants for each risk region, but that this number cannot exceed 20. The challenge is to find the “at most 20” causative variants amongst the thousands of genotyped or imputed variants in each locus. *ε_i_* is assumed to be normally distributed with mean 0 and variance 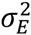.

#### Prior distributions

Following Sorensen and Gianola (2002), the values of *t* and 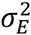 are fixed at 0 and 1, respectively. The individual liabilities, *y_i_*, are assumed to be normally distributed

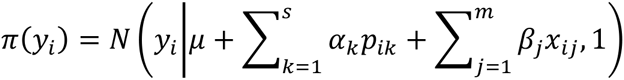

The population mean, *μ*, and effects of the PCs capturing population stratification are assumed to follow uniform distributions (*π*(*μ*) *∝* 1; *π*(*α_k_*) *∝* 1). Following Fang et al. (2012), the prior distribution of the effect of variant *j* from *m*, *β_j_*, is assumed to follow a double-exponential distribution:

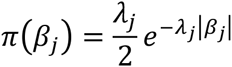

which is factorized in three sub-priors: (i) normally distributed 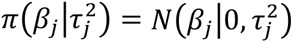; (ii) exponentially distributed 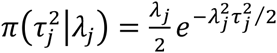; (iii) gamma distributed 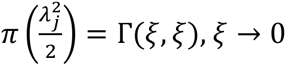.

#### Posterior distributions for Gibbs sampling

Effects *β_j_* are sampled from normal distributions with mean

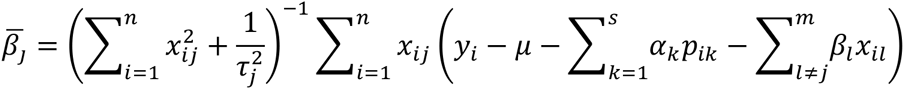

and variance

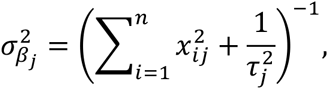

in which *n* is the total number of analyzed individuals (cases + controls). 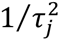 are sampled from inverse Gaussian distributions

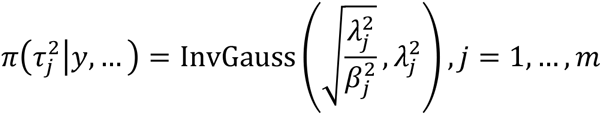

The hyper-parameters 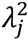 are sampled from gamma distributions

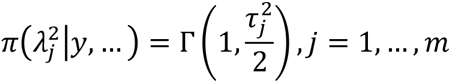

PC effects, *α_k_*, are sampled from normal distributions with mean

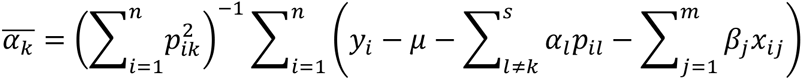

and variance

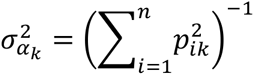

The population mean, *μ*, is sampled from a normal distribution with mean

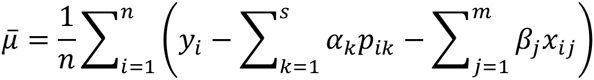

and variance 1/*n*.

For affected individuals (*γ_i_* = 1), the liabilities, *y_i_*, are sampled from the truncated normal distributions (such that *y_i_* > *t*) with density

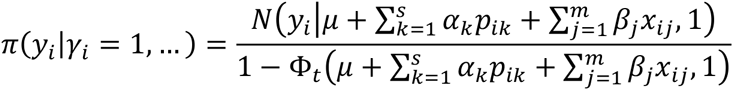

For unaffected individuals (*γ*_1_ = 0), the liabilities, *y_i_*, are sampled from the truncated normal distributions (such that *y_i_* ≤ *t*) with density

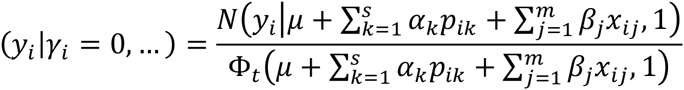

In these, 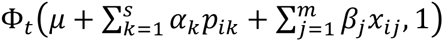 corresponds to the cumulative density from – ∞ to *t*.

#### Variant sampling using the Metropolis-Hastings algorithm

We first hierarchically cluster variants that are in high LD using (1-r^2^) as distance measure and the “single linkage” approach implemented with the R “hclust” package. By doing so variants in distinct clusters will never have r^2^ > C, yet variants within clusters may have r^2^ < C. We tested C values of 0.9 and 0.5. The *m* (=20) variants to include in the model are sampled such that each cluster can only be represented by one variant. At each round of the MCMC chain, we sequentially attempt to swap each of the *m* variants in the model with a better one, to ultimately find the best overall combination of variants. The substitute variants are selected 50% of the time from variants from the same cluster, and 50% of the time from variants of unrepresented clusters. In other words, this means that the MCMC chain spends halve of its time searching for the best possible variants within clusters, and halve of its time for the best possible clusters. When a substitute variant is selected, the probability to “accept” it is

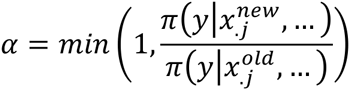

where

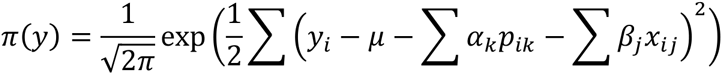

and 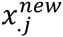 and 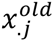 correspond to the genotype indicator variables for the “new” and “old” positions, respectively.

#### Implementation of the MCMC chain

We initiate the chain by assigning values randomly to all variables (within their legal boundaries). We then sequentially update the position of the *m* variants by either choosing a variant in another, unrepresented cluster (50%) or in the same cluster (50%), using the M-H algorithm described above. The likelihood of the new proposition is computed with the parameter values of the previous cycle. Corresponding 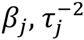 and *λ_j_* are updated by 50 rounds of Gibbs sampling. After each round of update of the *m* variants, we update *μ*, the *a_k_*‘s and *y_i_*‘s by one round of Gibbs sampling. The complete process was repeated 500,000 (simulated data) or 1 million times (real data). The first 100,000 (simulated data) or 500,000 cycles (real data) were used as burn-in and ignored when compiling the summary statistics.

#### Summarizing the results

We computed posterior probabilities (PP) for clusters as well as individual variants from the proportion of MCMC cycles in which they were included in the model. Within clusters, we defined “credible sets” of markers by ranking them on PP and considering the minimal set of markers that would jointly account for 95% of the PP of the cluster.

Clusters with posterior probability ≥ 0.50 were retained for further validation by fitting their corresponding lead SNPs jointly in a logistic regression model. Clusters exceeding the set significance threshold were considered to be positive. We used thresholds of 10^-4^, 10^-6^ and 10^-8^ that might be considered as locus-specific, multi-locus (~100 loci; cfr. Huang et al., 2016) and genome-wide thresholds.

### Forward Selection

The performance of BayesFM was compared with that of a standard forward selection approach implemented by logistic regression in R. Significance thresholds were the same as defined above (10^-4^, 10^-6^ and 10^-8^). We built credible sets associated with selected “lead” (*l*) variants by computing the PP for all *n* variants in high LD (r^2^ > C, as defined above) with *l*. The PP probability of variant *j* of *n* was computed as:

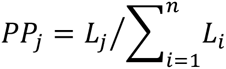

where *L_j_* is the maximum likelihood of the data considering variant *j* in the model. Likelihoods were computed using the glm() function (binomial family, logit link function, and logLik) in R. Credible sets (associated with a given lead variant) corresponded to the smallest set of variants that would jointly account for 95% of the PP. Variants included in a credible set were ignored when pursuing the forward selection.

## Datasets

### Simulated data

We took advantage of the Immunochip dataset of the International IBD Genetics Consortium (IIBDGC) and Multiple Sclerosis Genetics Consortium (IMSGC), consisting of 18,967 Crohn’s disease cases, 14,628 ulcerative colitis cases, and 34,257 controls, all of European ancestry. We randomly selected a genomic region corresponding to a GWAS-identified risk locus for Inflammatory Bowel Disease (chr5: 40,286,967–40.818,088) with 2,978 markers either interrogated by the Immunochip (936), or imputed from the 1,000 Genomes project with quality score > 0.4 (2042) (Huang et al., 2016). Within this region, we randomly selected one (model I), three (model II) or five (model III) variants with MAF ≥ 0.01, to act as causative variants. The variance explained by the locus 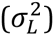 was set at 2% (see hereafter). In the cases with multiple causative variants, the proportion of the variance explained by the distinct causative variants was set at 4/7, 2/7 and 1/7 (three variants, model II) or 16/31, 8/31, 4/31, 2/31, 1/31 (five variants, model III).

The effect of a causative variant *j* (*β_j_*) was assumed to be additive and to have numerical value 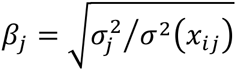, as 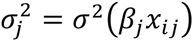. The value of 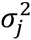, the variance due to variant *j*, was set as described above, i.e. 2% (model I) or the corresponding fraction of 2% (models II & III). The value of 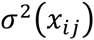, where *x_ij_* is – as before - the genotype dosage if individual *i* for variant *j*, was computed from the Immunochip data. The procedure described thus far does not account for the LD that may exist between the multiple causative variants in models II&III, which may cause 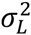 to deviate from 2%. Indeed,

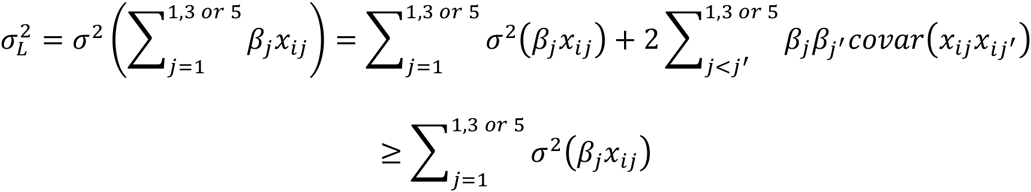

Thus, we rescaled the effects *β_j_* to 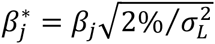 Substituting 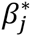 for *β_j_* in the previous equation indeed gives 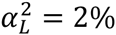.

To generate a simulated case-control cohort we sampled 1 million individuals from the Immunochip dataset with replacement and without discrimination of real case-control status. For each of these, we generated a liability, *y_i_*, as (i) the sum of the genotype effects at the 1, 3 or 5 causative variants: 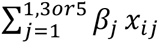, plus (ii) a residual effect, *ε_i_*, drawn from a normal distribution with mean 0 and variance of 1. Thus the variance explained by the locus as a fraction of the total liability variance is in fact 0.02/(1 + 0.02) « 0.02. Assuming an incidence of the disease of 1/400, we kept the 2,500 individuals with highest liability as case cohort. We randomly sampled 2,500 individuals from the remaining 1,000,000-2,500 = 997,500 individuals to serve as controls. We generated 100 such simulated case-control datasets to compare the performance of BayesFM with that of a more standard forward selection procedure. When analyzing the simulated datasets, we did not fit PC in the statistical models.

### Real data

For the analysis of real data, we likewise took advantage of the Immunochip dataset of the International IBD Genetics Consortium (IIBDGC) and Multiple Sclerosis Genetics Consortium (IMSGC), consisting of 18,967 Crohn’s disease cases, and 34,257 controls, all of European ancestry. We selected the genomic region corresponding to the GWAS-identified risk locus for Inflammatory Bowel Disease encompassing the *NOD2* gene (chr16: 50,692,364-50,847,022) with 1,048 markers either interrogated by the Immunochip (283), or imputed from the 1,000 Genomes project with quality score > 0.4 (765) (Huang et al., 2016). The analysis was restricted to Crohn’s disease.

## Results

### Simulated data

As expected, the True Positive Rate (TPR, i.e. the proportion of true positive signals amongst the total number of true signals (true positives plus false negatives)) was decreasing for both methods (BayesFM and FS) with increasing significance threshold and decreasing variance accounted for by the considered variant (Table 1 and Supplemental Table 1). Mapping resolution (defined as the size of the credible sets) was comparable between BayesFM and FS and only very mildly affected by significance threshold and variance explained. As expected, it decreased with LD threshold used to define clusters/credible sets (average number of variants per cluster/credible set of ~25 (r^2^=0.9) versus ~30 (r^2^=0.5)). Otherwise, LD threshold (r^2^=0.5 or 0.9) had only very modest effects on TPR.

For given thresholds and variance explained, the TPR tended to be slightly better for BayesFM than for FS, especially at the higher significance thresholds, but the differences were modest (Table 1 and Supplemental Table 1). However, when considering models II and III, characterized by multiple causative variants, the False Discovery Rate (FDR, i.e. the proportion of false positives amongst the total number of positive signals (true positives and false positives)) was considerably higher for FS than for BayesFM (Table 1 and Supplemental Table 1). Thus, while BayesFM and FS appear to have comparable sensitivity, BayesFM outperforms FS in generating a smaller number of false positives in situations of allelic heterogeneity.

To examine whether BayesFM and FS might be complementary and might best be used in combination as done in Huang et al. (2016), we measured the TPR and FDR for a specific scenario (model III, log(1/p) threshold 8) considering (i) both approaches separately, (ii) overlapping findings, and (iii) approach-specific findings. As expected the TPR was highest when considering both approaches individually. For the examined scenario, the sensitivity measured by the TPR was 31% for BayesFM and 28% for FS. The corresponding FDRs were 16% for BayesFM and 24% for FS. Hence and as mentioned before, BayesFM outperformed FS especially with regards to specificity in this scenario of allelic heterogeneity. When only considering positive results found by both methods (overlapping findings), the sensitivity dropped only very slightly when compared to FS alone (TPR = 26%), while the specificity increased considerably especially when compared to FS (FDR = 12.5%). Considering BayesFM-specific findings in addition to the overlapping findings (BayesFM and FS) increased the yield of true positives by nearly 20%, with a still reasonable FDR of 28%. Considering FS-specific findings in addition to the overlapping finings would only increase the yield of true positives by 9%, with an abysmal FDR of 68% (Figure 1).

**Figure 1:**
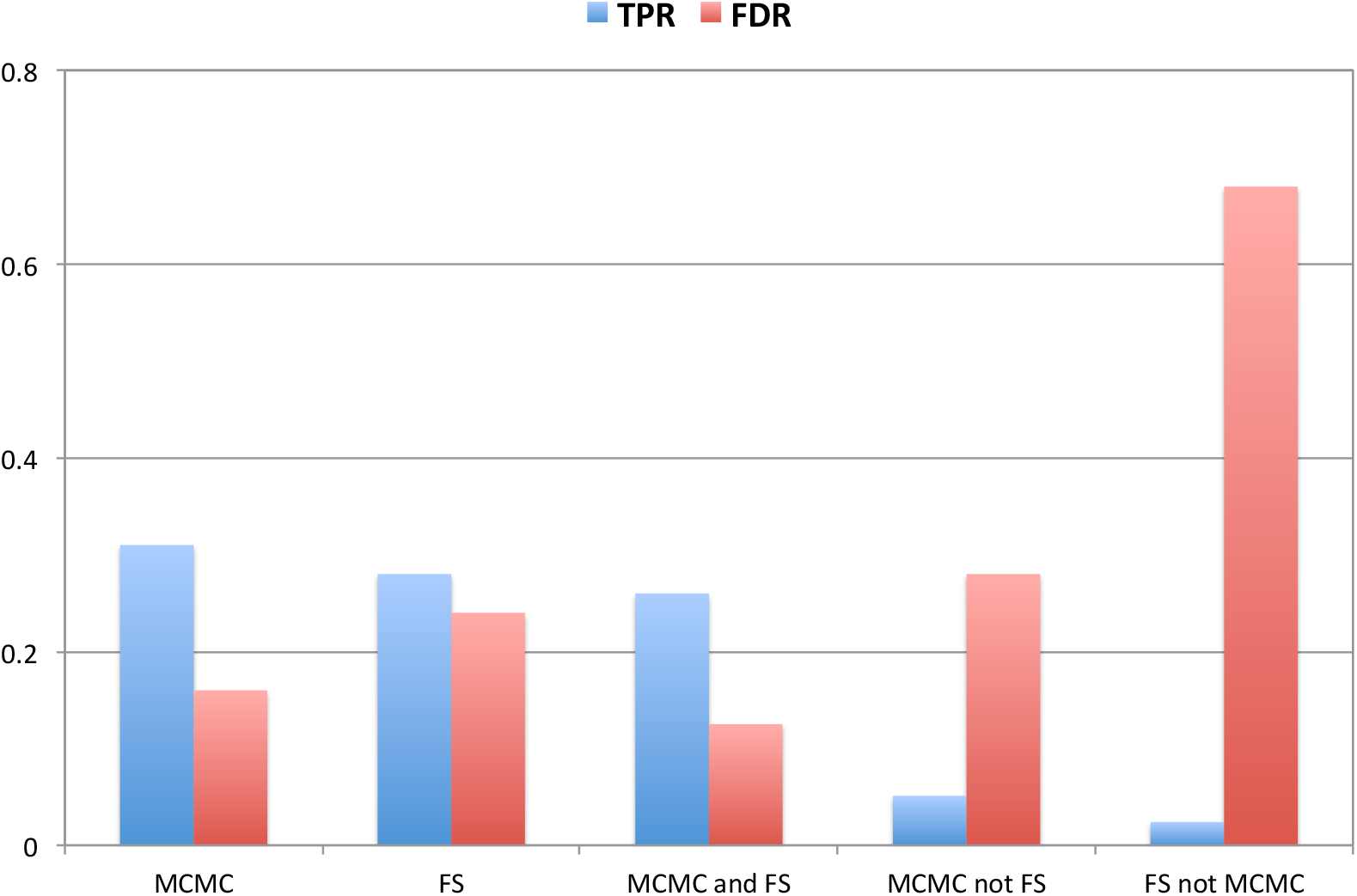
Simulated data. True Positive Rates (TPR) and False Discovery rates (FDR) obtained when (i) considering BayesFM (MCMC) and Forward Selection (FS) separately, (ii) when considering overlapping results (MCMC and FS), (iii) when considering method-specific results (MCMC not FS, and FS not MCMC). The scenario under consideration was model III (five causative variants) and a genome-wide significance threshold of log(1/p) = 8.

There are at least two scenarios in which BayesFM is expected to beat Forward Selection. The first is a situation in which a passenger variant is in LD with two or more causative variants and single-handedly accounts for a higher proportion of the variance than any of the causative variants alone. Standard Forward Selection will then irreversibly include it in the model, which may subsequently preclude the actual causative variants from entering it. An example of such a scenario, previously referred to as “ghost” effect, is illustrated in Figure 2A, and Table 2. It generates both false positives and false negatives with Forward Selection. A second scenario where BayesFM is expected to outperform Forward Selection is when two causative variants are in LD, and the risk alleles are in repulsion hence neutralizing each other effects. An example of such a situation is shown in Figure 2B and Table 2.

**Figure 2:**
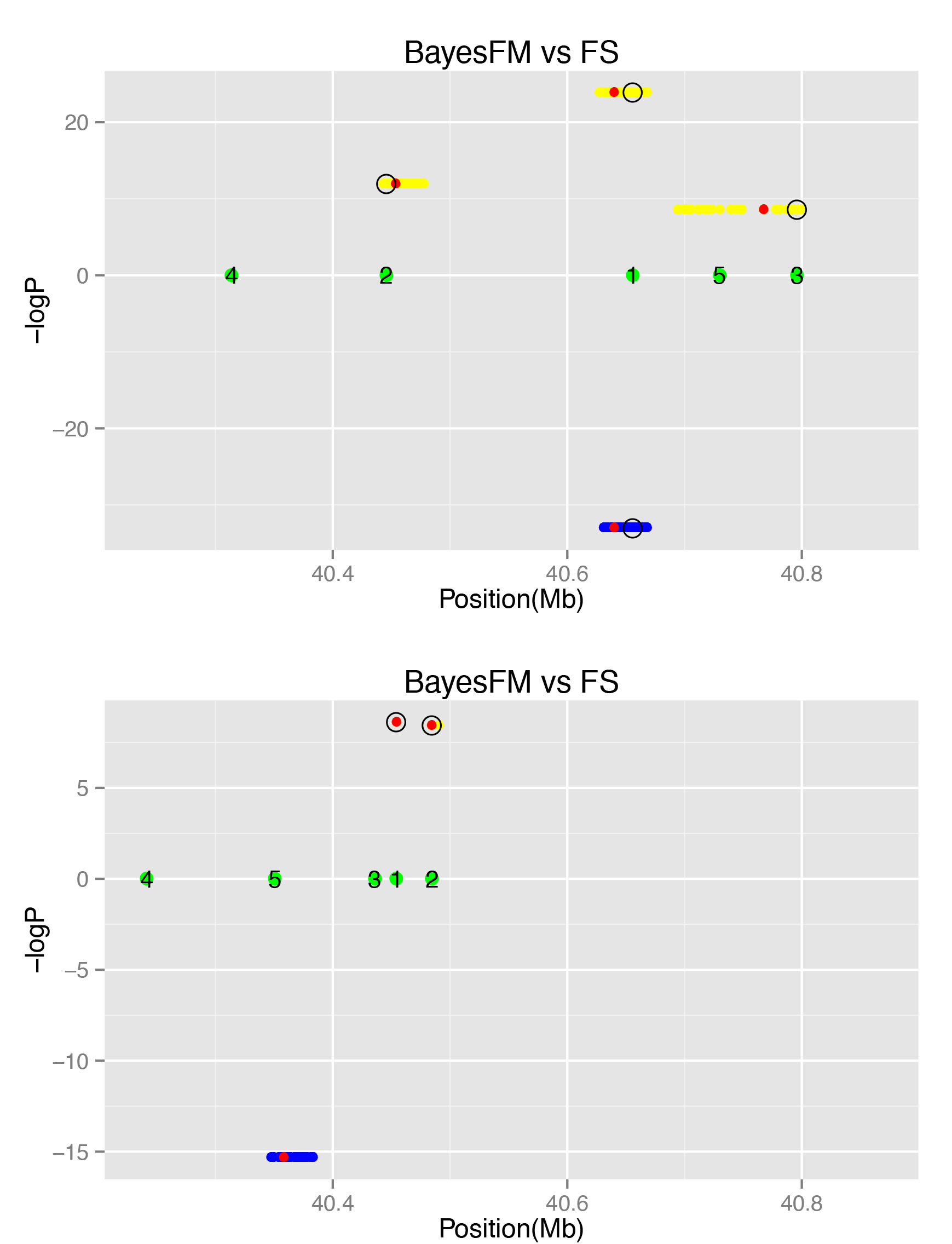
Simulated data. Statistical significance of disease association for credible sets of variants selected (PP ≥ 50%) using BayesFM (log(1/p), positive values) or Forward Selection (-log(1/p), negative values), estimated by multivariate logistic regression. The positions of the five simulated causative variants (model III) are shown by the numbered green dots. The yellow and blue dots mark the position of the variants in the credible sets identified by BayesFM and Forward Selection, respectively. The red dots mark the positions of the lead variants in the corresponding credible sets. Causative variants within credible sets are circled. **A.** Example of a “ghost QTL” effect. Forward Selection erroneously identifies a cluster of passenger variants that is in LD with multiple causative variants thereby single-handedly achieving a higher significance than any of the causative variants. It is therefore erroneously and irreversibly introduced into the forward selection model. BayesFM avoids this trap and correctly identifies at least causative variants 1 and 2. **B.** Example of two causative variants in LD with an excess of haplotypes with risk alleles in repulsion. By modeling them simultaneously, BayesFM uncovers risk alleles 2 and 3. By modeling them sequentially, Forward Selection misses both 2 and 3 as they neutralize each other’s effects.

### Real data

We then examined the results obtained with BayesFM on the *NOD2* locus (chr16: 50,692,364–50,847,022), a well-established risk locus for Crohn’s disease. We analyzed the dataset of the IIBDGC described in Huang et al. (2016) and comprising 18,967 Crohn’s disease cases and 34,257 matched controls. Table 3 summarizes the results that were obtained using either r^2^ > 0.9 or r^2^ > 0.5 as LD threshold to define clusters of variants (see M&M). For both analyses, we report the clusters with PP > 0.50. Thirteen such signals were obtained with r^2^ > 0.9, and fourteen with r^2^ > 0.5. The average number of variants per signal was 1.15 with r^2^ > 0.9 and 3.14 with r^2^ > 0.5. Single variant resolution was obtained for 11/13 signals with r^2^ > 0.9 and 8/14 signals with r^2^ > 0.5, highlighting the remarkable resolving power that can be achieved for at least some loci with BayesFM. The log(1/p) value obtained by fitting the lead variant (with highest PP) of each signal in a multivariate logistic regression, exceeded 6 for 10/13 signals with r^2^ > 0.9 and 10/14 signals with r^2^ > 0.9. The lowest log(1/p) value was 3.33 for the signal that was detected using r^2^ > 0.5 only. It was > 4 for all others.

Using very stringent criteria, nine independent signals were retained for the same locus and reported in Huang et al. (2016)(Table 3). All but one of these were detected by BayesFM, whether using r^2^ >0.9 or 0.5. These included four signals corresponding to single variant resolution of non-synonymous (NS) variants in *NOD2* (fs1007insC, R702W, G908R, N289S, all with PP~1), and one signal with two-variant resolution corresponding to two non-synonymous variants each (V793M and S431L; resolved to V793M by BayesFM when setting r^2^ >0.9). The remarkable enrichment in NS variants testifies of the specificity of the fine-mapping methods utilized in Huang et al. (2016) including BayesFM. The signal that remained undetected by BayesFM was characterized by a PP of 0.24 with r^2^>0.9 and 0.20 with r^2^>0.5. It is worth noting that the corresponding lead variant *(rs104895467)* has a Phastcons conservation score of 1, increasing the likelihood of it being a genuine causative variant.

Amongst the six additional putative signals detected by BayesFM when applying more lenient thresholds, three were characterized by a NS *NOD2* lead variants (A585T, R676C and A891D). This remarkable enrichment in NS variants strongly suggests that at least some of these signals are true.

Two of the detected signals are characterized by very common risk alleles (31% and 61% in cases, respectively). Both of these signals are characterized by fairly large clusters when using either FS or BayesFM (r^2^>0.5), indicating that many variants are in high LD with the corresponding causative variants. Four of the signals are characterized by low frequency risk alleles (1% < frequency < 5% in controls) and include the well-known *fs1007incC, R702W, G908R* and *rs72796367* variants. The remaining signals correspond to rare risk alleles with frequencies below 1% in controls.

## Discussion

Identifying causative variants in GWAS-defined risk loci is important in order to gain a better understanding of the molecular mechanisms underlying inherited disease predisposition, including the identification of the causative genes. A number of fine-mapping strategies have been explored to achieve this goal using association information in case-control cohorts. These include Bayesian approaches to define credible sets that are likely to contain the causative variants (f.i. Wellcome Trust Case Control Consortium et al., 2012; van de Bunt et al., 2015). However, the corresponding methods make the unlikely assumption of allelic homogeneity at the considered loci, i.e. the occurrence of single causative variants only. Searching for multiple independent causative variants in a given locus – a more realistic scenario - is most often conducted using variations of stepwise forward selection approaches. In these approaches - if deemed significant - the strongest signal is sequentially added as covariate to the model. Two of the methods utilized in Huang et al. (2016) are advanced Bayesian versions of this approach. It is relatively easy to imagine scenarios in which these forward selection approaches may either miss true signals or incorporate non-causative variants into the model (see results section on simulated data). Alternative Bayesian variable selection approaches, combined with Monte Carlo techniques, have been devised to overcome these limitations. In the field of fine-mapping, these include BimBam (Servin & Stephens, 2007), GUESSFM (Wallace et al., 2015), and BayesFM presented in this manuscript. Simulations (including in this study) indicate that they are generally superior to the other approaches. In our simulations, BayesFM appeared to have improved sensitivity and specificity when compared to a standard implementation of forward selection. Our analyses confirm the benefits of selecting signals that are detected by forward selection *and* BayesFM, the strategy followed in part by Huang et al. (2016). It considerably improves specificity with limited impact on sensitivity. We nevertheless note that the signals detected by BayesFM only (and not FS) appear to be characterized by an acceptable specificity, while FS-only signals were in essence not trustworthy (Figure 1).

Ideally, fine-mapping should be done with complete sequence information in the utilized case-controls. While this may become possible in the near future as sequencing costs continue to diminish, present studies typically augment genotyping data from SNP arrays with imputation using the for instance the 1,000 Genomes Project data as reference (f.i. Huang et al., 2016). The imputation accuracy varies between variants and is typically inferior for low frequency variants. This is very likely affecting the precision of fine-mapping and very difficult to overcome by. An accurate evaluation of the impact of imputation accuracy on the outcome of fine-mapping, including with Bayesian model selection approaches such as BayesFM is needed.

In its present version, BayesFM assumes that the identified risk variants operate “additively”, i.e. we ignore the possibility of dominance within variants and epistatic interaction between variants. BimBam allows modeling of dominance effect within variants (Servin & Stephens, 2007). The impact of this simplification on power and accuracy remains to be determined. Modeling these higher order effects implies the estimation of additional parameters. As a consequence, impact on power and accuracy is likely to be a function of sample size.

The results of our simulations (data simulated under a simplified additive model) indicate that fine-mapping results need to be considered with caution. FDR was > 10% even when considering signals detected both by FS and BayesFM, and this is likely to be an underestimate. Nevertheless, applying FS-type methods in conjunction with BayesFM on a large case-control cohort for Inflammatory Bowel Disease lead to the fine-mapping of 42 signals with resolution < 5 variants. The causality of the corresponding variants, especially when non-coding, can now be tested directly by CRISPR-CAS9 technology in cell culture systems.

## Acknowledgments

This work was funded by the Welbio CAUSIBD, Belspo BeMGI and ARC IBD@ULg grants. We are grateful to the IIBDGC for allowing the use of the consortium genotype data, and Hailang Huang, Luke Jostins, Mark Daly and Jeff Barrett for fruitful discussions.

## Software availability

BayesFM can be downloaded from https://sourceforge.net/projects/bayesfm-mcmc-v1-0

## References

Fang M, Jiang D, Yang R, Fu W, Pu L, Gao H, Wang G, Yu L. 2012. Improved LASSO priors for shrinkage quantitative trait loci mapping, Theor Appl Genet 124: 1315–1324.

Huang H, Fang, Jostins L, et al. 2016. Association mapping of inflammatory bowel disease loci to single variant resolution. Nature, under revision.

Jostins L, Ripke S, Weersma R, Duerr R, McGovern D, et al. 2012. Host-microbe interactions have shaped the genetic architecture of inflammatory bowel disease, Nature 491: 119–124.

Servin B, Stephens M. 2007. Imputation-based analysis of association studies: candidate regions and quantitative traits. PLoS Genet 3: e114.

Sorensen D & Gianola D. 2002. Likelihood, Bayesian and MCMC Methods in Quantitative Genetics. Springer.

Van de Bunt M, Cortes A, IGAS Consortium, Brown MA, Morris AP, McCarthy MI. 2015. Evaluating the performance of fine-mapping strategies at common variant GWAS loci. PLoS Genet 11: e1005535.

Wallace C, Cutler AJ, Pontikos N, Pekalski ML, Burren OS, Cooper JD, Garcia AR, Ferreira RC, Guo H, Walker NM, Smyth DJ, Rich SS, Onengut-Gumuscu S, Sawcer SJ, Ban M, Richardson S, Toff JA, Wicker LS. 2015. Dissection of a complex disease susceptibility region using a Bayesian stochastic search approach to fine mapping. PLoS Genet 11: e1005272.

Wellcome Trust Case Control Consortium, Maller JB, McVean G, Bymes J, Vukcevic D, et al. 2012. Bayesian refinement of association signals for 14 loci in 3 common diseases. Nat Genet 44:1294–1301.

Welter D, MacArthur J, Morales J, Burdett T, Hall P, Junkins H, Klemm A, Flicek P, Manolio T, Hindorff L, and Parkinson H. 2014. The NHGRI GWAS Catalog, a curated resource of SNP-trait associations. Nucleic Acids Research 42 (Database issue): D1001–D1006.

